# Motor cortex retains and reorients neural dynamics during motor imagery

**DOI:** 10.1101/2023.01.17.524394

**Authors:** Brian M. Dekleva, Raeed H. Chowdhury, Aaron P. Batista, Steven M. Chase, Byron M. Yu, Michael L. Boninger, Jennifer L. Collinger

## Abstract

The most prominent role of motor cortex is generating patterns of neural activity that lead to movement, but it is also active when we simply imagine movements in the absence of actual motor output. Despite decades of behavioral and imaging studies, it is unknown how the specific activity patterns and temporal dynamics within motor cortex during covert motor imagery relate to those during motor execution. Here we recorded intracortical activity from the motor cortex of two people with residual wrist function following incomplete spinal cord injury as they performed both actual and imagined isometric wrist extensions. We found that we could decompose the population-level activity into orthogonal subspaces such that one set of components was similarly active during both action and imagery, and others were only active during a single task type—action or imagery. Although they inhabited orthogonal neural dimensions, the action-unique and imagery-unique subspaces contained a strikingly similar set of dynamical features. Our results suggest that during motor imagery, motor cortex maintains the same overall population dynamics as during execution by recreating the missing components related to motor output and/or feedback within a unique imagery-only subspace.

## Introduction

As people prepare to execute a skilled action, they often pause beforehand to mentally rehearse and visualize it. For example, a tennis player might imagine hitting an upcoming serve or a pianist might imagine playing a difficult sequence prior to performance. This type of covert motor imagery is constrained to the same performance limits as exist for overt execution. One study showed that the speed with which people were able to imagine performing a sequence of finger movements was limited to their actual overt performance (Sirigu et al., 1996). Motor imagery is similarly impacted by neurologic impairment; a lesion to motor cortex leads to an equal slowing of both executed and imagined movements (Sirigu et al., 1995). Conversely, imagery-based practice can improve actual motor function, in some instances offering a comparable performance benefit as standard overt training (Clark, 1960; Frank et al., 2014; Ladda et al., 2021; Yue & Cole, 1992). This tight coupling between imagery and actual motor function suggests similar central mechanisms, so that information and experience gleaned from one modality can usefully inform the other.

Primary motor cortex, known mainly for its role in the generation of volitional movement, is also active during covert motor imagery (Jeannerod, 2006; Kilteni et al., 2018; Rastogi et al., 2020; Vargas-Irwin et al., 2018). In fact, many movement-related brain areas are also active during the mental rehearsal of imagined movements. Premotor and supplementary motor cortices (Cisek & Kalaska, 2004; Stephan et al., 1995), anterior cingulate areas (Stephan et al., 1995) and parietal areas (Aflalo et al., 2015; Stephan et al., 1995) all display modulated activity during covert motor imagery. Despite the clear link between imagery and action, we know little about how cortical population activity differs (if at all) between the two. We hypothesize that the structure of imagery activity parallels that of movement preparation. These two concepts are certainly distinct; movement preparation involves “readying” for an imminent overt action, while motor imagery is the covert rehearsal of a complete action. However, during both imagery and preparation, the motor system faces a similar objective: to engage in movement-related processing while avoiding the activation of descending control pathways. In the case of movement preparation, the cortical implementation appears to be well explained by the coordination of a small number of correlation patterns within the neural population (Elsayed et al., 2016; Kaufman et al., 2014). Within this framework, activity in motor and/or premotor cortices activate orthogonal neural subspaces during preparation and execution. This orthogonality allows preparatory activity to evolve while avoiding the dimensions that would engage descending pathways and cause overt movement.

Given that imagery by definition does not involve overt movement, it seems reasonable to assume that imagery activity in motor cortices, like preparatory activity, avoids dimensions responsible for downstream control. However, this is possible under several distinct organizations of the population activity. One possibility is that imagery could exist only in a subset of the neural dimensions underlying action activity, or another possibility is that it could engage completely separate neural dimensions. Additionally, if imagery does engage some (or entirely) unique neural dimensions, the specific temporal responses within those dimensions, i.e. neural dynamics, could take any form.

Here we use an isometric wrist extension task to examine the relationship between imagery and action in motor cortex. Two participants with tetraplegia due to spinal cord injury participated in the study. Despite having no hand or lower extremity function, both retained residual proximal arm and wrist extension control. We recorded from intracortical microelectrode arrays implanted in the hand and arm areas of motor cortex as they performed either real or imagined isometric wrist extensions to achieve low or high force targets. After reducing the recorded population activity to a low dimensional manifold, we found that it contained three distinct subspaces: (1) a shared space, in which responses were nearly identical during action and imagery, (2) a unique action subspace that contained significant modulation only during actual force production, and (3) a unique imagery subspace that contained significant modulation only during imagery. Strikingly, we found that the neural dynamics within the unique imagery subspace during imagery closely resembled those observed in the unique action subspace during execution. From this, we conclude that motor cortex maintains the same overall neural dynamics during imagery as it contains during overt action. However, since the population activity must avoid output dimensions (and lacks feedback activity) during imagery, cortex instead recapitulates output/feedback responses within an orthogonal, imagery-unique subspace. We propose that the retention of overall neural dynamics structure during imagery provides the motor system with a useful proxy for overt practice.

## Results

### Motor cortex is active during both actual and imagined isometric wrist extensions

We asked participants to perform isometric wrist extensions within an immobile frame affixed with a force sensor to control the height of a line trace displayed on a monitor in front of them (Figure 1a). During “action” trials, they observed a horizontal bar indicating the required force (either low: ~5N or high: ~40N) and then were required to apply the appropriate force such that the line trace matched the vertical position of the bar. During “imagery” trials, the participants were required to imagine producing those same forces without actually doing so. On imagery trials, the line trace automatically increased to the target force and then returned to zero. For each session, we collected alternating twelve-trial blocks of action and imagery, resulting in thirty-six total trials of action and thirty-six trials of imagery. We then removed trials with force profiles that deviated significantly from the average (see *Experimental Setup*), resulting in approximately 31 ± 4 action trials and 28 ± 4 imagery trials per session (six sessions for P2 and three sessions for P3).

**Figure 1.**
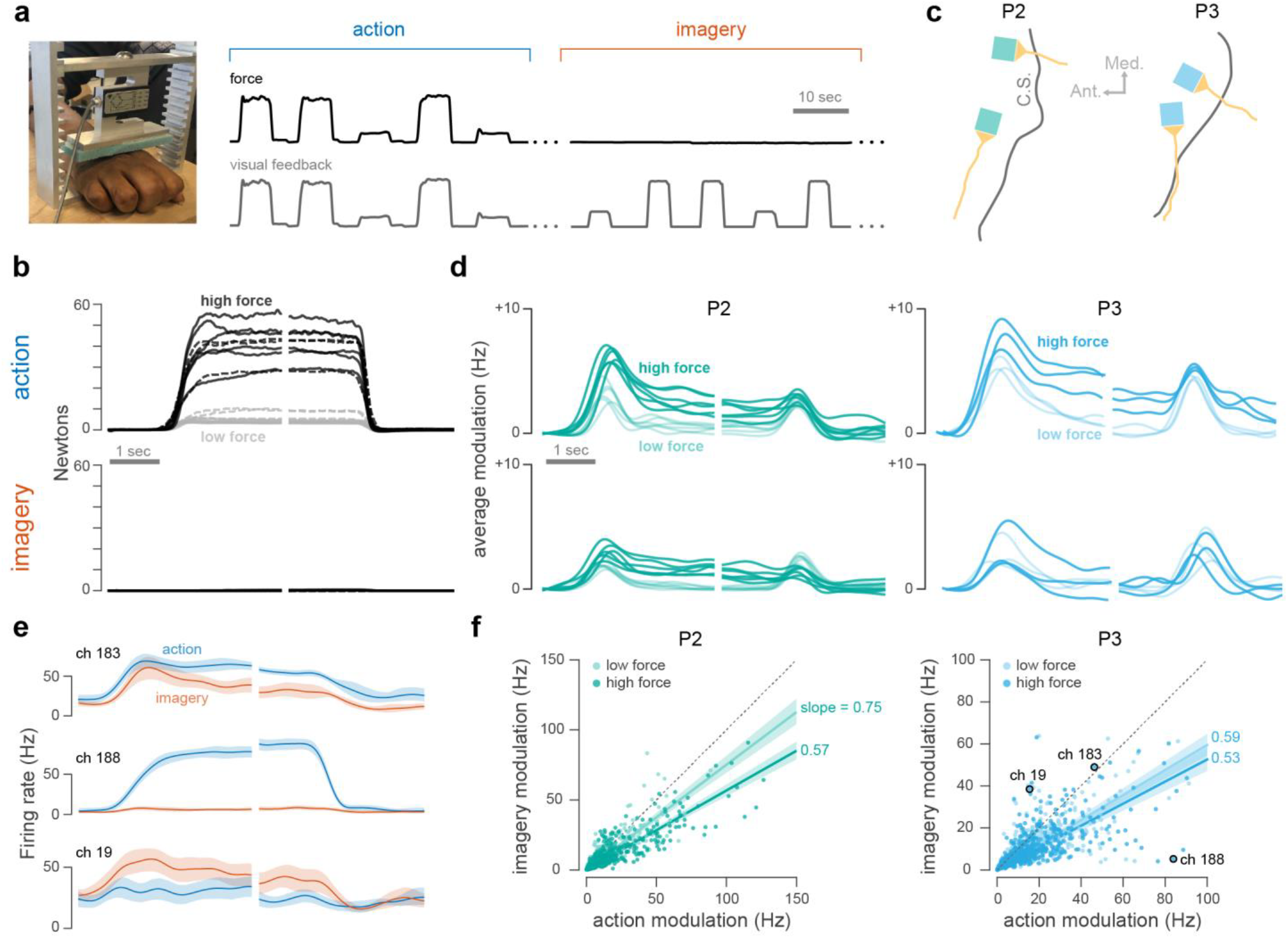
Motor cortex is active during both imagery and action. **(a)** Participants placed their hands on a board beneath a load cell and produced either real or imagined wrist extension forces. For all trials, they received visual feedback of either their actual produced force or an automated proxy. **(b)** Average low force (gray) and high force (black) traces for all sessions (P2: solid, P3: dashed) during action trials (top) and imagery trials (bottom). **(c)** Locations of microelectrode arrays implanted in the motor cortices. C.S. = central sulcus. **(d)** Average population firing rate modulation in motor cortex (M1) during action trials (top) and imagery trials (bottom). Each trace corresponds to a single session. Lighter traces represent low force trials and darker traces represent high force trials. **(e)** Average activity for three example channels (P3) during high-force action (blue) and high-force imagery (red) trials. **(f)** Maximum modulation during low force (light) and high force (dark) action and imagery for all recorded channels and all sessions (left: P2, right: P3).

Both participants successfully achieved and maintained the requested force targets during action trials (Figure 1b, top), and produced no significant force during imagery trials (Figure 1b, bottom). Throughout the experimental sessions, we recorded activity from the hand and arm areas of motor cortex (Figure 1c). Despite the stark difference in force output between action and imagery trials, we observed relatively subtle change in the overall population-wide firing rates between the two task types (Figure 1d, top vs. bottom). However, individual channels displayed a wide variety of response types, including a mix of preferential activation during action or imagery (Figure 1e). Some channels displayed similar modulation during both action and imagery (e.g., Figure 1e, channel 183), while others appeared uniquely activate during only one task type (e.g., Figure 1e, channels 188 and 19). For each channel, we calculated the maximum modulation during action and imagery by calculating the difference between the 5^th^ percentile and 95^th^ percentile firing rates for each task type (imagery or action) and force level. We found that for P2, the average low-force imagery modulation was 75 ± 6% that of low-force action modulation. High-force imagery modulation was 57 ± 4% that of action (Figure 1f, left). For P3, low-force imagery modulation was 59 ± 5% that of action and high-force imagery modulation was 53 ± 6% that of action.

### Neural latent space contains distinct action and imagery subspaces

As a first step towards characterizing the differences between the neural representations of action and imagery, we asked whether the population activity could be separated into unique dimensions containing only action or imagery activity. For simple motor behaviors, the measured dimensionality of motor cortical activity is typically far lower than the number of recorded neurons (Elsayed & Cunningham, 2017; Gallego et al., 2017, 2020; Remington et al., 2018; Sadtler et al., 2014; Shenoy et al., 2013). Thus, as an initial step, we reduced the activity from our recorded populations (176 channels for P2, 192 channels for P3) to a twenty-dimensional latent space. To do this, we first performed factor analysis (FA) separately on action and imagery trials to obtain two ten-dimensional latent spaces and then combined them into a single twenty-dimensional space that spanned both initial ten-dimensional spaces (see *Methods: Dimensionality reduction* for details). The initial ten-dimensional FA space computed from action trials accounted for 70% of action variance (P2: 76 ± 3%, P3: 59 ± 2%), but only 51% of imagery variance (P2: 59 ± 6%, P3: 36 ± 4%). The initial ten-dimensional FA space computed from imagery trials accounted for 69% of imagery variance (P2: 75 ± 5%, P3: 58 ± 7%), but only 54% of action variance (P2: 63 ± 5%, P3: 37 ± 5%). The combined twenty-dimensional space accounted for 78% of action variance (P2: 83 ± 3%, P3: 69 ± 2%) and 75% of imagery variance (P2: 80 ± 3%, P3: 65 ± 6%). Thus, by combining low-dimensional subspaces identified independently for each task, we were able to obtain a single subspace that captured the majority of meaningful activity for both action and imagery.

Each dimension of the resulting twenty-dimensional latent space captured non-zero variance during both action and imagery (Figure 2a, top). To separate activity that was unique to action and imagery, we performed an iterative procedure in which we identified—one at a time—dimensions that contained significant variance for only a single task type (see *Methods: Subspace separation*). The identified dimensions were constrained to be orthonormal, such that the latent space remained unchanged (i.e. the percentages in Figure 2a sum to 100%). Thus, the final transformation simply provides a different view of the same underlying latent space, such that activity clusters into discrete subspaces with unique task-related variance characteristics (Figure 2a, bottom). Specifically, we found three relevant orthogonal subspaces: an action-unique subspace containing significant variance only during the action task, an imagery-unique subspace containing significant variance only during the imagery task, and a shared subspace containing both action and imagery variance. We also isolated a null subspace that contained little activity for either task, which likely resulted from an overestimation of the total neural dimensionality.

**Figure 2.**
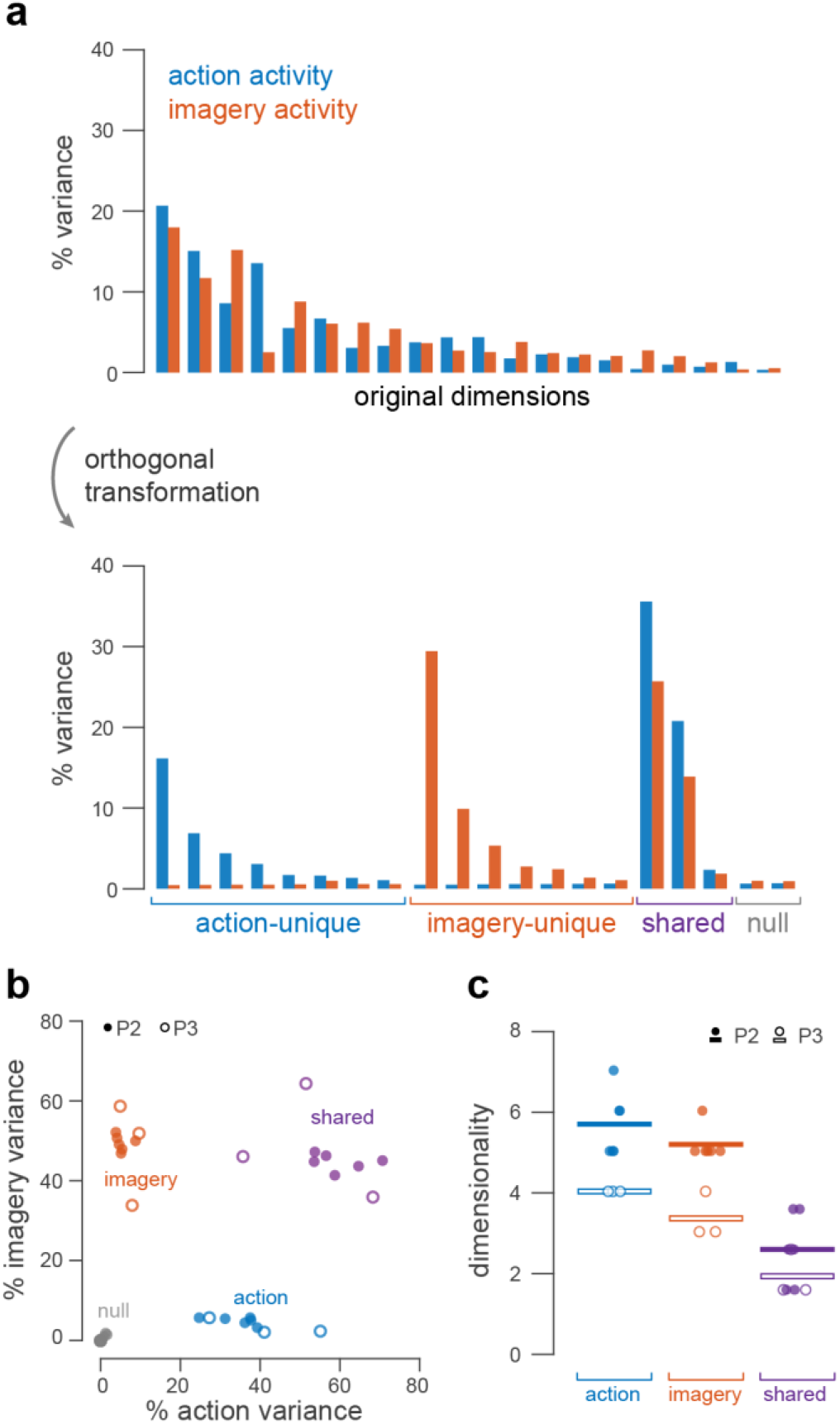
Population activity contains distinct action, imagery, and shared subspaces. **(a)** Percent of total latent variance explained by each dimension for an example session (P2), for both action (blue) and imagery (red) tasks. Top shows distribution for original factor dimensions, and bottom for the same space following an orthogonal rotation to separate action and imagery variances. **(b)** Percent action and imagery variance captured by the action, imagery, shared, and null subspaces. Solid circles correspond to sessions from P2, open circles to sessions from P3. **(c)** Dimensionality (number of dimensions that explain at least 1% total variance) of action, imagery, and shared subspaces. Solid circles correspond to individual sessions from P2, open circles to sessions from P3. Solid and open bars represent cross-session averages.

We found that for both the action and imagery tasks, the total population variance was distributed between the shared subspace and relevant unique subspace. After splitting the population activity into action, imagery, and shared subspaces, we calculated the explained variance of each (Figure 2b). For either task, the explained variance of the opposite subspace (e.g. action subspace during the imagery task) never exceeded 10% of the task-specific variance. For the action task, the action subspace explained approximately 38% of the variance (P2: 35 ± 6%, P3: 41 ± 14%), while the shared subspace explained 56% (P2: 60 ± 7%, P3: 52 ± 16%). For the imagery task, the imagery subspace explained approximately 49% of the variance (P2: 50 ± 2%, P3: 48 ± 13%), while the shared subspace explained 47% (P2: 45 ± 2%, P3: 49 ± 14%).

In addition to total variance explained, we also examined the dimensionality of each subspace (Figure 2c). To do this, we performed principal components analysis (PCA) on the activity within each subspace and counted the number of dimensions that accounted for more than one percent of the total variance. We found that, on average, the dimensionalities of the action and imagery subspaces were similar (P2: 5-6D, P3: 3-4D) and larger than the dimensionality of the shared subspace (P2: 3D, P3: 2-3D).

### Temporal components in the shared subspace are equivalent during action and imagery

The previous analysis of the neural latent space revealed three separate subspaces in which neural activity evolves, depending on whether the subject exerted force or imagined exerting force. Next, we turn to an examination of how neural activity evolves in time within each of these subspaces. We first used each participant’s trial-averaged neural responses to align each of the three subspaces across sessions (see *Cross-session subspace alignment*). We then compared trial-averaged activity during the action and imagery tasks within the aligned shared subspace.

Projecting the trial-averaged activity from both tasks into the shared subspace revealed a striking similarity in the temporal profiles between action and imagery for each shared dimension (Figure 3a). The action-imagery correlation values in Figure 3a correspond to the separate cross-session correlations along the three axes displayed. However, these axes (and corresponding correlations values) represent just one view of the three-dimensional shared subspace activity. To better assess the correlation irrespective of the selected coordinate frame, we performed a Monte Carlo method (see *Monte Carlo sampling*) in which we sampled 10000 random unit vectors from the shared subspace (Figure 3c). On each draw, we computed the correlation between the action and imagery activity along that dimension. Across all sampled dimensions, we found median correlations of 0.95 (P2) and 0.90 (P3), indicating that the shared subspace activity was universally well matched between the two task types.

**Figure 3.**
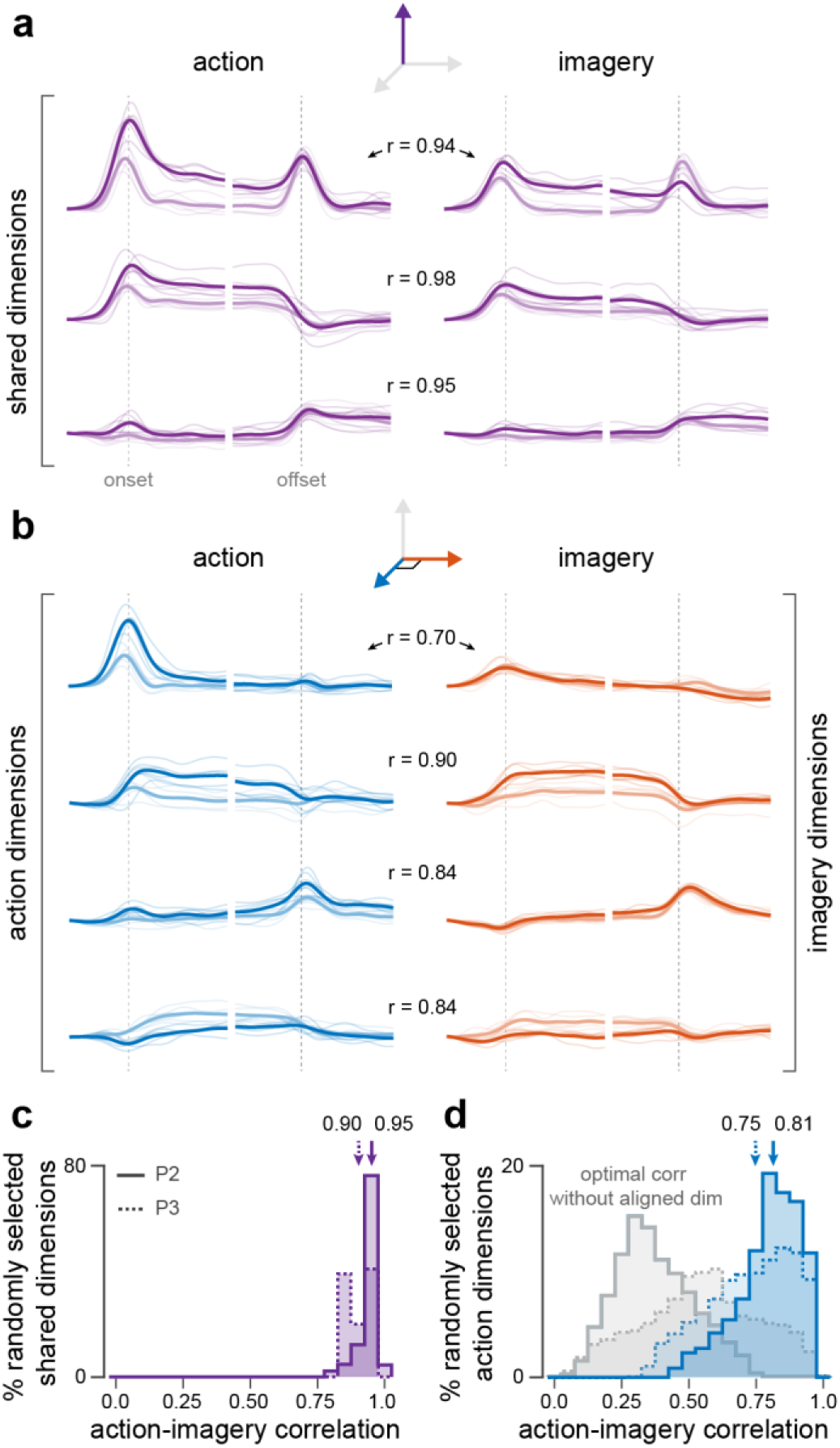
Temporal components of action and imagery are similar for both shared and unique subspaces. **(a)** Action activity (left) and imagery activity (right) projected onto the shared dimensions (P2). Thin traces correspond to individual sessions, thick lines to cross-session averages. Light and dark traces represent low and high force averages, respectively. **(b)** Action activity projected onto action dimensions (left) and imagery activity projected onto imagery dimensions (right), following an alignment procedure (see *Action-Imagery subspace alignment*). Correlation coefficients describe the correlation between action activity projected on action dimensions and imagery activity projected onto imagery dimensions. **(c)** Action-imagery correlations across 10,000 randomly selected individual dimensions from the shared subspace. Arrows show median correlations for each participant. **(d)** Same as in (c), but calculated between action dimensions and aligned imagery dimensions. Gray distributions represent the maximal possible correlations if the corresponding aligned imagery dimension is removed on each random draw.

### Temporal components in the imagery subspace match those in the action subspace

The comparison of temporal components between the action and imagery subspaces is less straightforward than for the shared subspace. By construction, the action and imagery subspaces are orthogonal to each other and only contain meaningful activity during their respective tasks, making it futile to compare activity along a single dimension within these subspaces. Instead, we found a rotation of the imagery subspace axes that matched the multidimensional responses observed during imagery to the action subspace responses observed during action. (see *Action-Imagery subspace alignment*). Following this alignment procedure, we found that the action and imagery spaces comprised a similar set of temporal components (Figure 3b), despite existing in orthogonal subspaces of neural activity.

We quantified the overall similarity between the multidimensional action and imagery responses by calculating correlations between the action activity on randomly chosen dimensions in the action subspace and the imagery activity on the corresponding (aligned) imagery dimensions (Figure 3d, blue). Across all randomly sampled dimensions, we found median correlations of 0.81 (P2) and 0.75 (P3). To provide context for these values, we also found, for each randomly selected action dimension, the maximally correlated imagery dimension that was not the aligned dimension (Figure 3d, gray). These secondary dimensions represent the correlation with “the next best dimension” and thus reflect the degree of triviality in the original alignment. Temporal components reflecting nonspecific task timing, for example, could be fairly ubiquitous and correlates might exist to a similar degree on multiple dimensions. However, we found that activity on aligned imagery dimensions was uniquely correlated with the corresponding action dimension activity; removing the aligned imagery dimension consistently reduced the maximum possible correlation that could be achieved from all other dimensions (P2: p<0.0001, P3: p<0.0001, Wilcoxon signed-rank test). Thus, the set of temporal components found in the action-unique subspace are reoriented into an orthogonal imagery-unique subspace during mental imagery.

### Action- and imagery-unique activity is integral to the dynamical structure of M1

There is ample evidence that—at least for simple, practiced tasks—activity in M1 can be largely described by simple linear dynamics (Churchland et al., 2012; Perich et al., 2020; Shenoy et al., 2013). One consequence of this dynamical structure is the presence of low neural tangling in the population activity (Russo et al., 2018). That is, each neural state is predictive of a unique subsequent state; there are no points from which activity wildly diverges for different task phases. Our decomposition of the neural activity into shared and unique dimensions allows us to examine how these separate subspaces contribute to the untangled, dynamical nature of M1 activity.

We first calculated tangling of the neural response in the full D-dimensional latent space, comprising both the shared and unique subspaces. To then determine the extent to which this tangling depended on temporal components from the shared subspace, we performed a simple dropout analysis, visualized in Figure 4a (see *Neural tangling* for details). Briefly, we found a single dimension from the shared subspace that, when removed, resulted in the highest tangling in the remaining (D−1)-dimensional space. We performed the same procedure for the unique subspaces, again removing a single action-unique (or imagery-unique) dimension that maximized tangling. We found that for both action and imagery, removing shared dimensions led to a non-significant ~32% increase in tangling (Figure 4b,c; *action*: P2: +47 ± 46% (p=0.07, paired KS test), P3: +15 ± 24% (p=0.81), *imagery*: P2: +26 ± 24% (p=0.43), P3: +34 ± 16% (p=0.08)). However, removing unique dimensions led to a large (~265%) and significant increase in tangling (Figure 4b,c; *action*: P2: +300 ± 203% (p=0.00015, paired KS test), P3: +314 ± 118% (p=0.0013), *imagery*: P2: +244 ± 161% (p=0.00092), P3: +190 ± 90% (p=0.0013)). We then directly compared tangling between the two dropout conditions (shared dimension removed versus unique dimension removed) and show that across conditions and participants, removing dimensions from the unique subspaces lead to consistently higher tangling than removing dimensions from the shared subspace (Figure 4d,e). This result suggests that activity within the unique subspaces has a greater contribution to maintaining low tangling in the population response than activity within the shared subspace.

**Figure 4.**
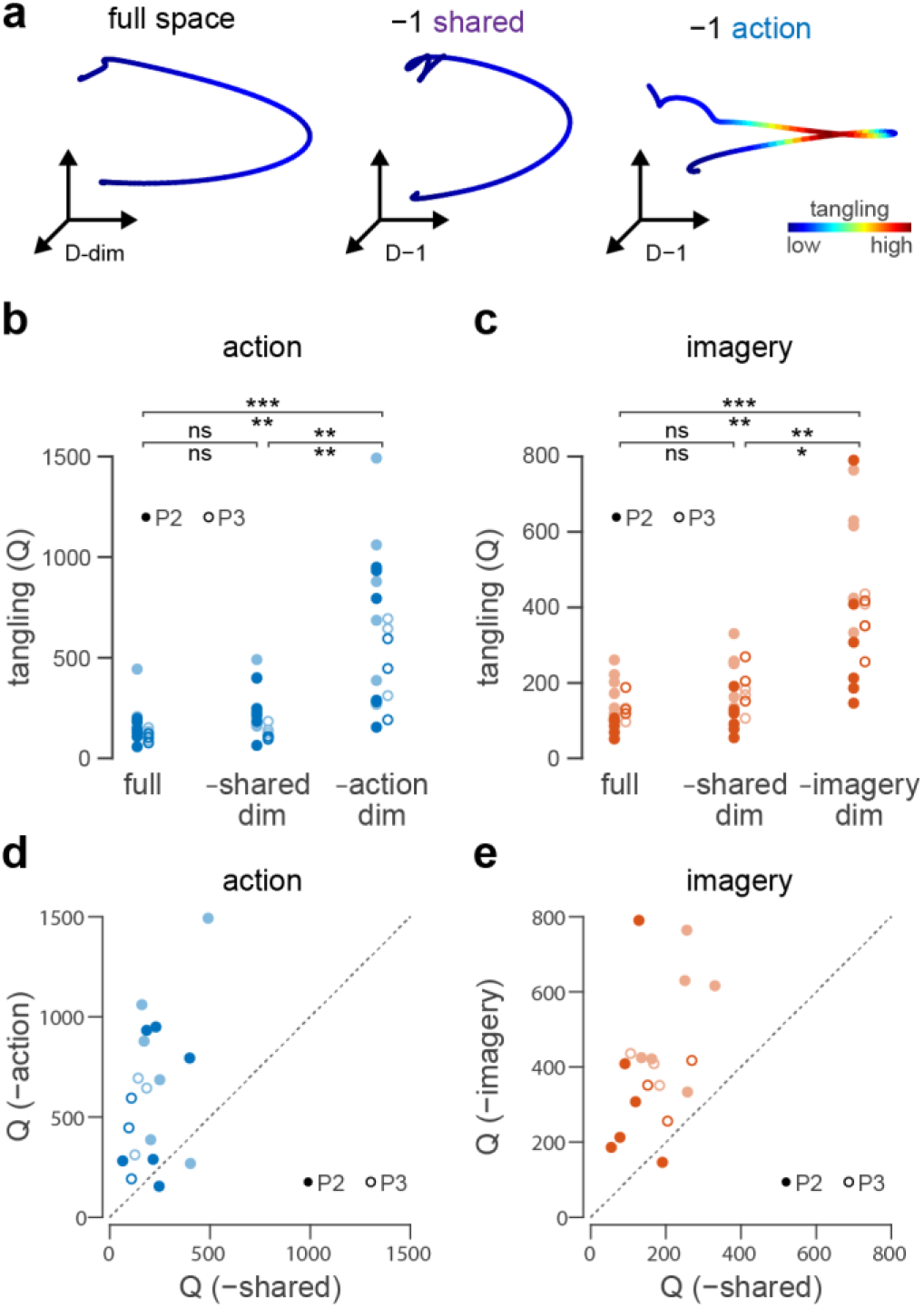
Action-unique and imagery-unique subspaces are responsible for maintaining low neural tangling **(a)** *Left*: Projection of the total D-dimensional latent activity during the onset period the high force condition, colored according to tangling (blue=low tangling, red=high tangling). *Middle*: Projection showing tangling in the (D−1)-dimensional latent space after removing a shared dimension to maximize tangling. *Right*: Projection showing tangling in the (D−1)-dimensional latent space after removing an action dimension to maximize tangling. **(b)** Maximum tangling for trial-averaged responses of each session and force condition. Light and dark points indicate low and high forces, respectively. Significance between groups indicated for P2 (above line) and P3 (below line) based on two-sample KS test **(c)** Same as in (b) for imagery conditions **(d)** Comparison of maximum tangling between (D−1)-dimensional spaces, in which a shared (x-axis) or action (y-axis) dimension has been removed. **(e)** Same as in (d) for imagery trials, comparing tangling after removing either a shared dimension or an imagery dimension.

## Discussion

In this study, we examined the relationship between population activity in motor cortex (M1) during isometric force production and corresponding covert motor imagery. We found that the low-dimensional manifold activity comprised three orthogonal subspaces: a shared subspace, a unique action subspace, and a unique imagery subspace. Activity within the shared subspace accounted for approximately half of the total variance and was nearly identical during both action and imagery. Activity in the action and imagery subspaces, though constructed from completely orthogonal correlation patterns, also contained well-matched sets of temporal responses. This action-unique and imagery-unique activity appeared essential for maintaining a simple dynamical structure within M1. Removing individual dimensions of action-unique or imagery-unique activity led to a dramatic increase in tangling, whereas removing dimensions of shared activity had almost no impact.

Because it is only active during action (and not during motor imagery), we hypothesize that the action-unique subspace that we identified is directly involved in generating motor output and possibly receiving sensory feedback. During imagery, output dimensions must be avoided and there is no incoming somatosensory feedback. In theory, motor cortex could satisfy the constraint of avoiding output dimensions by restricting imagery activity to a lower-dimensional subspace, such that there existed only a shared subspace and an action-unique subspace. However, we instead found that imagery engaged an additional, separate subspace, which contained temporal components equivalent to those in the action-unique (putative “output”) subspace. This suggests that covertly imagining an action does not simply suppress output activity, but instead rotates it into dimensions that do not generate muscle activity (Figure 5). There are multiple potential explanations for why the act of motor imagery should involve creating “dummy” output-like responses in motor cortex. One reason might be that during imagery, cortex is practicing to generate output commands, even though those commands do not actually make it downstream. Even for this one-dimensional task, the action-unique subspace contained between four and seven dimensions (Figure 2c), which suggests an output space that is more complex than the eventual muscle activity. In addition to muscle-like responses, (Figure 3b, second component) the action-unique subspace also contains transient responses (Figure 3b, first component), which could reflect indirect control through subcortical areas. There is evidence that downstream motor structures are able to integrate brief, transient activity from cortex to generate sustained muscle output (Shalit et al., 2012; Albert et al., 2019). The ability to practice producing this multidimensional control signal within a motor-output-null space before generating the actual output commands might explain why mental rehearsal improves subsequent performance on overt motor tasks (Clark, 1960; Frank et al., 2014; Ryan & Simons, 1982; Schack et al., 2014; Sheahan et al., 2018; Yue & Cole, 1992).

**Figure 5.**
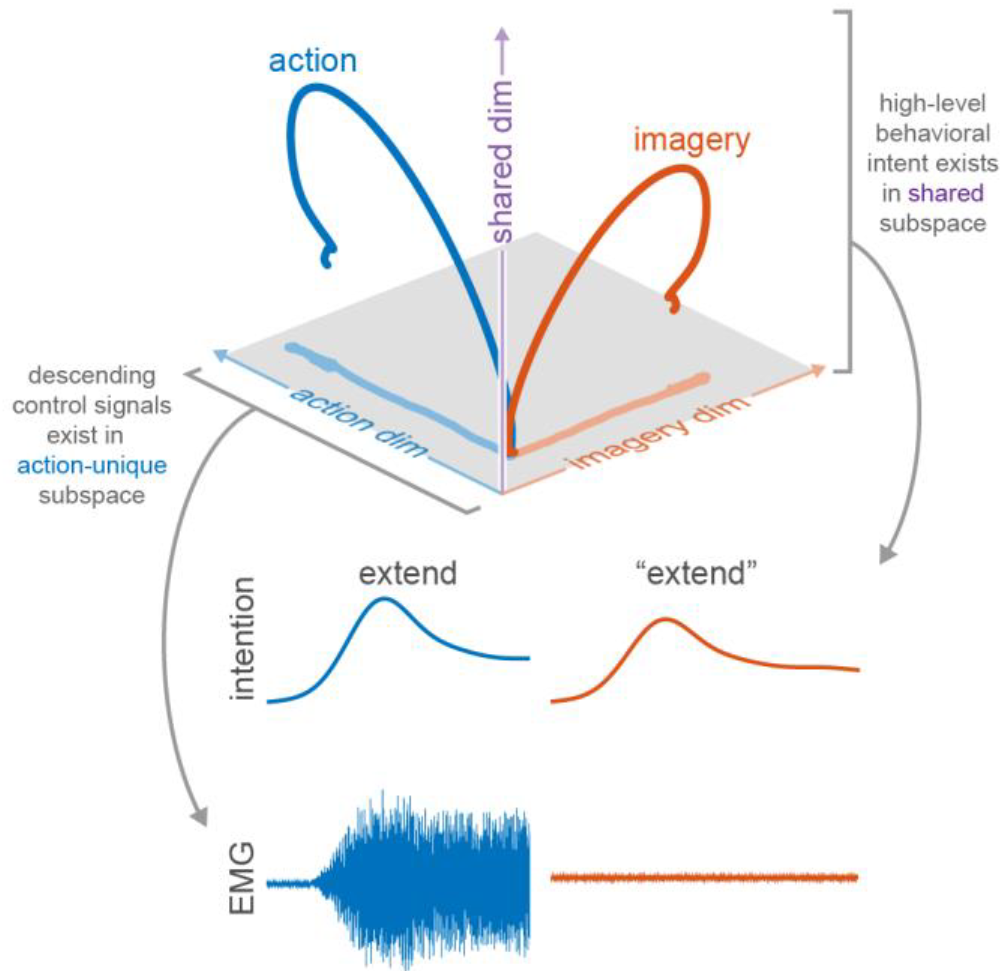
Hypothesized population-level architecture of motor cortex that enables both action and imagery.

A separate possible reason for the existence of output-like components during imagery is that they are necessary for maintaining the dynamical structure of the entire motor cortical ensemble. There is ample evidence that for simple, predominantly feedforward behaviors, motor cortex activity resembles a nearly autonomous dynamical system (Churchland et al., 2012; Perich et al., 2020; Shenoy et al., 2013). The multidimensional population response unfolds predictably from an initial neural state, often dictated by preparatory activity in premotor cortex (Elsayed et al., 2016), presumably reflecting intrinsic M1 or broader synaptic connectivity. From our results, it appears that output-related activity (i.e. activity in the action-unique subspace) is largely responsible for maintaining this untangled, simple dynamical structure in motor cortex. Removing just a single dimension of action-unique or imagery-unique activity caused a significant increase in tangling, while removing shared activity had almost no discernible effect (Figure 4). Therefore, suppressing action-unique activity entirely during imagery would lead to a complete collapse of the dynamic structure in motor cortex. However, recapitulating those components in an orthogonal subspace suppresses output while preserving its dynamical properties, which might be important for stabilizing the behavior of the broader sensorimotor network across different volitional states.

The orthogonality between action and imagery subspaces presumably functions similarly to the orthogonality observed between movement preparation and movement execution (Elsayed et al., 2016; Kaufman et al., 2014). For both preparation and imagery, restricting the population activity to uniquely non-output dimensions prevents unwanted movement. However, the two processes are distinct in their dynamic relationship to the action. If enough time is allowed, preparatory activity in premotor areas appears to settle into a static neural state (Cisek & Kalaska, 2005). From the dynamical systems perspective, this represents an initial set point, which dictates how the subsequent multidimensional response will unfold during movement execution (Churchland et al., 2010). The process of motor imagery, on the other hand, does not reflect an imminent preparation for movement, but rather rehearsal of the entire action. The placements of the microelectrode arrays in this study were too close to the central sulcus to observe strong preparatory responses, so we do not know how preparation for imagery might relate to preparation for action. Uncovering the full population-level organization of preparation-, imagery-, and action-related activity could help elucidate the processes by which cortex uses both overt and covert processes to improve motor skill.

The concept of covert motor imagery also invokes the related function of action observation. When people or animals observe others performing a motor action, it also engages motor-related brain areas in a way similar to self-initiated movement (Dushanova & Donoghue, 2010; Hari et al., 1998, 1998; Holmes et al., 2006; Jiang et al., 2020; Muthukumaraswamy & Johnson, 2004; Papadourakis & Raos, 2019; Rizzolatti et al., 1996; Stefan et al., 2005; Tkach et al., 2007). This correlation between observation and action exists even in the activity of individual cortical neurons (Cisek & Kalaska, 2004; Dushanova & Donoghue, 2010; Mazurek et al., 2018; Tkach et al., 2007; Vigneswaran et al., 2013), which supports the notion of a “mirror neuron” network through which the motor system can presumably learn new skills by observing others (Rizzolatti et al., 2001). It is tempting to assume that observation and imagery/rehearsal are equivalent processes, and that observing an action triggers a person (or animal) to imagine performing the action themselves. Since the vast majority of intracortical observation-based experiments are performed with monkeys, it is often impossible to resolve the degree to which the subject is actively involved in motor imagery. Recent work in humans with tetraplegia found that observation and imagery are actually not equivalent (Rastogi et al., 2020; Vargas-Irwin et al., 2018), and that it is possible to distinguish those two volitional states from population-level activity in motor cortex. Our observation of orthogonal subspaces containing action-unique and imagery-unique activity mirrors results from Jiang et al. (2020), who observed separate subspaces containing action-unique and observation-unique activity, suggesting that although observation and imagery can be considered distinct volitional states, they might employ similar population-level mechanisms (i.e. orthogonal subspaces for output and non-output conditions). However, because the task used in our study was isometric, we could not include a meaningful observation-only condition. In the future, including kinematic limb movements would help identify the degree of overlap (common versus unique subspaces) across a larger range of volitional conditions, including observation, imagery, action, and even sleep (Rubin et al., 2022).

While the task-dependent nature of the action and imagery subspaces provides clear insight into their functional roles (e.g. the action-unique subspace includes output-related activity), interpretation of the shared subspace is more difficult. Activity in this subspace was nearly identical for both tasks, suggesting that it represents some sort of higher-level task objective. The separation of force levels within the shared subspace argues against the interpretation that it is highly nonspecific and reflects broad subject state processes like arousal or engagement (Hennig et al., 2021). Instead, it appears to contain information related to the specific task goal (i.e. force level). We speculate that activity in the shared subspace corresponds to goal-oriented or “task-intention” signals from higher order brain areas.

From an ethological perspective, the ability to imagine movements is only useful if it can meaningfully inform or assist overt motor control. Here we show that activity in motor cortex during imagery is essentially dynamically equivalent to activity during action. One subset of the multidimensional activity is shared across tasks, occupying the same subspace during both action and imagery. The remaining subset of activity evolves in orthogonal subspaces for action and imagery, but with strikingly similar dynamics. Overall, our results illuminate the population-level architecture by which motor cortex is able to engage in both movement generation and covert motor imagery.

## Methods

### Participants

Two participants (P2 and P3) took part in this study. P2 is a 35-year-old man with tetraplegia caused by C5 motor/C6 sensory ASIA B spinal cord injury. P3 is a 30-year-old man with tetraplegia caused by incomplete C6/C7 ASIA B spinal cord injury. Both participants retain some residual upper arm and wrist control, but no hand function.

Both participants had two microelectrode arrays (Blackrock Microsystems, Salt Lake City, UT) implanted in the hand and arm areas of motor cortex (P2: two 88-channel arrays, P3: two 96-channel arrays). They also had two 64-channel arrays implanted in somatosensory cortex (Flesher et al., 2016), which were not used for this study. Informed consent was obtained prior to performing any study-related procedures. Data collection for P2 occurred approximately five years post-implant, and collection for P3 approximately one year post-implant.

### Experimental Setup

For each experimental session, the participants placed their pronated right hands on a board on their lap. We then secured a load cell in a frame attached to the board, positioning it such that it made gentle contact with the top of the hand. At the start of each session, we asked participants to perform one maximal voluntary contraction (MVC). We then set initial low and high force targets based on the peak force observed during MVC (10% and 60%) and asked participants to practice by attempting each force level a few times. If the participants reported concern that the high force target was too high and he would be unable to perform the task without significant pain or fatigue, we lowered it until it reached a comfortable level. Once the force targets were set, we began the experiment, alternating blocks (12 trials each for P2, 10 trials each for P3) of action and imagery. Participant P2 performed three blocks of action and three blocks of imagery for all sessions. Participant P3 performed four blocks of each for sessions one and two, and five blocks for session three. On each block, we randomly interleaved low and high forces. Each session always began with action to provide a reference for the subsequent imagery. We excluded trials that contained force traces that significantly deviated from the cued profile. On average, we excluded 6 ± 2 action trials and 8 ± 4 imagery trials (due to non-zero force output) per session.

### Data acquisition

We collected neural data via digital NeuroPlex E headstages connected via fiber optic cable to two synced Neural Signal Processors (Blackrock Microsystems). The neural signals were filtered using a 4^th^ order 250 Hz high-pass filter, logged as threshold crossings (−4.5 RMS) and subsequently binned at 50 Hz. These binned counts were then square root transformed and convolved offline with a Gaussian kernel (σ = 200 ms) to provide a smoothed estimate of firing rate.

### Dimensionality reduction

We sought to reduce the dimensionality of the neural population recordings by projecting the activity into a low-dimensional space. Principal components analysis (PCA) or factor analysis (FA) are often used to perform such a reduction. However, we also wanted to take into account any differences in mean firing rate and/or modulation between action and imagery; standard PCA or FA would tend to return leading dimensions that preferentially captured action-related variance, since the action task contained higher overall variance. To ensure that the low-dimensional space captured a similar percentage of variance for each task, we performed a modified FA procedure.

As a first step, we performed FA independently for the action and imagery tasks using z-scored rates. This provided two weight matrices, ***W**_action_* ∈ ℝ^*N*×10^ and ***W**_imagery_* ∈ ℝ^*N*×10^ (*N* = 176 for P2, *N* = 192 for P3) that described transformations for projecting channel activity to separate (but highly overlapping) 10-dimensional spaces. We then concatenated these two matrices into a new matrix ***W**_action+imagery_* ∈ ℝ^*N*×20^ and performed singular value decomposition,

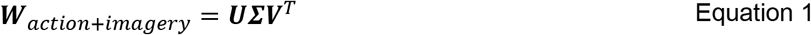

This procedure provided ***U*** ∈ ℝ^*N*×20^, an orthonormal basis spanning both ***W**_action_* and ***W**_imagery_*. The resulting twenty-dimensional space is almost certainly an overestimation of the actual dimensionality. However, this allowed for the possibility that imagery and action activity existed in entirely orthogonal subspaces. We projected data from both tasks into this space by first normalizing the rate of each channel by the sum of its individual variances from both tasks and multiplying the resulting normalized rates by ***U***.

We calculated the percent variance explained by each subspace ***W*** of the full *N*-dimensional mean-centered neural data ***Z***:

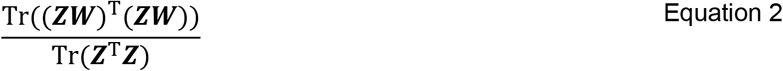

### Subspace separation

We aimed to divide, if possible, the low-dimensional manifold into orthogonal subspaces for which the activity exhibited unique task-related variance characteristics. Essentially, we wanted to identify subspaces that contained wholly task-specific variance (only variance during action or imagery). To achieve this, we implemented a modified version of a subspace identification method introduced by Jiang et al. (2020). The original method by Jiang et al. identifies a task-unique subspace e.g. ***Q**_task1_* by optimizing:

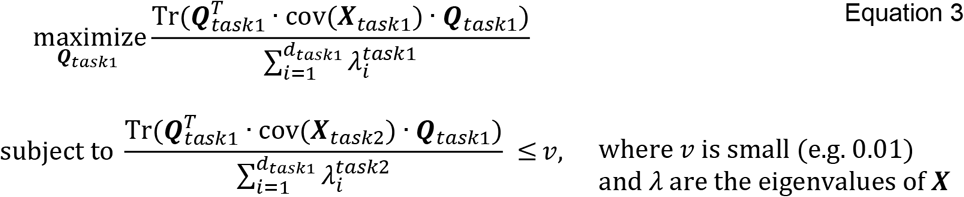

The resulting subspace ***Q**_task1_* contains the maximum variance from task 1 under the constraint that the captured variance from task 2 must not exceed a set (small) amount. For our specific purposes, this exact formulation has two drawbacks: (1) the dimensionality of ***Q*** must be defined *a priori*, and (2) the resulting ***Q**_task1_* and ***Q**_task2_* are not orthogonal to each other. To address these limitations, we created a modified, iterative implementation to identify unique subspaces one dimension at a time.

1. Initialize ***Q**_action_* and ***Q**_imagery_* to be empty.
2. For latent factor data ***X**_action_* and ***X**_imagery_* (both with current dimensionality *d*), separately perform the optimization from Equation 3 to find (if possible) *d*-dimensional vectors ***v**_action_* and ***v**_imagery_* (***v*** takes the place of ***Q*** in Equation 3).
3. If:
  a. both ***v**_action_* and ***v**_imagery_* satisfy the low cross-variance constraint, then select the one that explains the most within-task variance. This vector defines a new dimension of ***Q**_action_ or **Q**_imagery_*.
  b. only one satisfies the constraint, it defines a new dimension of the associated subspace ***Q***.
  c. neither satisfies the constraint, stop
4. Reduce factor data to the (*d-1*)-dimensional space that is orthogonal to the current ***Q**_action_* and ***Q**_imagery_*
5. Repeat 2-4 until 3c.

The subspace remaining after the procedure terminates represents the shared space, since it does not include any dimensions for which variance from one task is separable from variance from the other task. We performed the optimization using the Manopt toolbox in Matlab (Boumal et al., 2014).

### Cross-session subspace alignment

We decomposed the low-dimensional factor space activity into action, imagery, and shared subspaces (as described in *Subspace separation*) for each session independently. Recording instabilities can change the activity observed on single channels across sessions (Perge et al., 2013), making it impossible to apply a single transformation (e.g. projection to low-dimensional subspaces) across multiple days. However, there is evidence that low-dimensional (manifold) representations of the population activity remain consistent for a single behavior (Gallego et al., 2020). Therefore, to combine datasets from different sessions, it is necessary to first align the low-dimensional spaces (Degenhart et al., 2020). We chose to align subspace activity (action, imagery, and shared) for each participant using Generalized Procrustes Analysis (GPA; Gower, 1975). GPA iterates to find a multidimensional response common to all datasets and returns the axis transformation (here we used only an orthonormal rotation with no scaling) necessary to align each dataset with that common response.

With the subspace responses aligned across sessions, we then found a compact representation of the underlying temporal components of each subspace using a varimax rotation. That is, we performed a varimax rotation on the baseline-centered multidimensional common response (identified by GPA) for each subspace. We defined “baseline” as the average response during a 200 ms window at the start of the trial period. We then ranked the varimax-rotated version of the multidimensional response by variance. As an orthogonal transformation, this varimax procedure did not change the underlying nature of the multidimensional responses, but rather highlighted the separable temporal features within each subspace. For example, the first three dimensions in the action subspace (Figure 3b) are readily interpretable as onset, sustained, and offset responses, respectively.

### Action-Imagery subspace alignment

Just as the low-dimensional subspace responses had to be aligned to make cross-session comparisons, so too did the action and imagery subspaces. We wanted to compare the similarity of the multidimensional temporal responses between the action and imagery subspaces. However, we could not simply compare, e.g. the first dimension of the action subspace with the first dimension of the imagery subspace since the two subspaces had undergone separate cross-session alignment procedures (see *Cross-session subspace alignment*).

To compare the imagery and action subspace responses, we found an orthonormal transformation, ***Z**_im-act_*, of the imagery subspace that maximized the sum of the squared covariance between action responses in the action subspace and imagery responses in the imagery subspace.

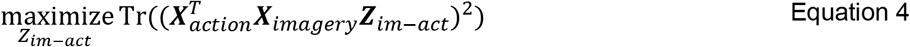

We performed this optimization using the Manopt toolbox in Matlab (Boumal et al., 2014). Unlike the more commonly used canonical correlation analysis for aligning temporal components (Gallego et al., 2020; Jude et al., 2022), this rotation is orthonormal and represents a middle ground between maximizing correlation and returning leading components that explain a large amount of within-subspace variance.

### Monte Carlo sampling

For the temporal response comparisons in Figure 3c,d we wanted to quantify the similarity of the multidimensional responses in a way that did not depend on the chosen coordinate frame. For example, we provide in Figure 3a,b the specific correlation values for the displayed dimensions, but an orthogonal rotation of the space—which preserves the actual multidimensional relationship—would result in a different set of correlations.

To provide a coordinate-frame-agnostic quantification of two multidimensional responses, we performed a Monte Carlo sampling-based procedure in which we calculated the correlation between responses on randomly selected dimensions.

For multidimensional data ***X**_1_* and ***X**_2_* (dimensionality *d*):

1. Generate a random *d*-dimensional unit vector ***u**_rand_*.
2. Project ***X**_1_* and ***X**_2_* onto ***u**_rand_* and calculate the correlation between the resulting projections

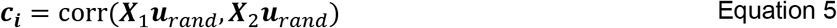
3. Repeat for 10,000 random vectors

The resulting distribution of correlations ***c*** provides an overall picture of the multidimensional correspondence between ***X**_1_* and ***X**_2_*.

### Action-Imagery correlation control distributions

The Monte Carlo sampling-based method provided a quantification of the overall correlation between responses in the action and imagery subspaces following the imagery-action alignment (*Action-Imagery subspace alignment*). However, we also wanted to include an additional reference distribution that would help provide context for the resulting distribution of correlations (Figure 3d). Ideally, we aimed to clarify whether the relatively high correlations observed between the action and imagery subspaces were *unique* to the specific alignment (indicating a true alignment of similar components), or if they simply reflected broad, nonspecific modulation throughout the multidimensional space as a result of e.g. task timing.

The core question that we addressed with our control procedure was: for any given dimension, what is the maximum possible correlation between action and imagery if we remove the aligned imagery dimension? To implement this, we performed an additional optimization on each draw of the Monte Carlo routine (*Monte Carlo sampling*).

1. Project imagery subspace activity ***X**_imagery_* into the (*d*-1)-dimensional space orthogonal to ***u**_rand_* (i.e. in the nullspace of ***u**_rand_*) to obtain ***X**_im-V-null_*.
2. Find unit vector ***m**_im-V-null_* that maximizes the correlation between ***X**_im-V-null_* and ***X**_action_**u**_rand_*

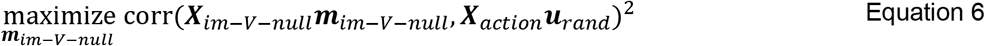
3. Save resulting (positive) correlation

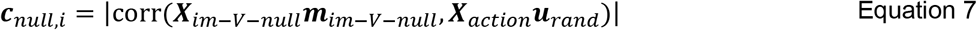

The distribution of ***c**_null_* is therefore a strong control, since (unlike ***c***) each element results from an independent optimization routine. However, even with that additional freedom, the values of ***c**_null_* were consistently lower than those in ***c***. This indicates that the temporal structure of action activity along any given action dimension is *uniquely* mirrored by imagery activity along the corresponding matched imagery dimension.

### Neural tangling

We used the metric of neural tangling (Russo et al., 2018) to examine the influence of each subspace (action, imagery, shared) on the overall dynamical nature of population M1 activity. This metric reflects the degree to which all states along a neural trajectory are predictive of subsequent neural states, which indicates the degree to which the activity follows predictable neural dynamics.

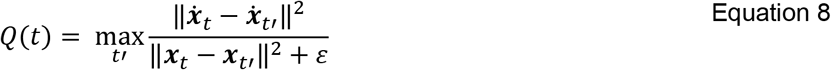

where ***x**_t_ and **ẋ**_t_* represent the neural state and derivative of the neural state at time *t*, and ε is a small value (one tenth of total variance of ***x***) to prevent a zero in the denominator.

For the dropout procedure used in Figure 4, we aimed to find a single dimension from either the shared, action-unique, or imagery-unique subspace that, when removed, resulted in the highest tangling in the remaining space. For example, to find the maximum tangling after removing a single action-unique dimension, we performed the following:

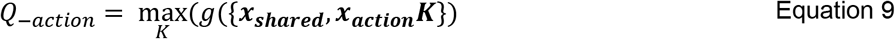

where

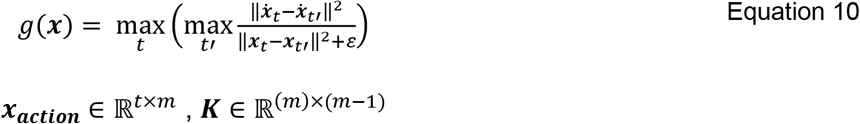

We performed this optimization using the Manopt toolbox in Matlab (Boumal et al., 2014). We also confirmed the results by performing a brute force search. To do this, we performed 10,000 iterations in which we selected a random vector from e.g. the action-unique subspace and calculated the tangling in the remaining space. The maximum tangling values over all random draws were always lower than the maximum tangling estimated via optimization.

## Acknowledgments

We would like to thank Nathan Copeland and Mr. Dom for their continued efforts and commitment to this study. We would also like to thank the research team, especially Debbie Harrington for regulatory management as well as Caroline Schoenewald, Jordyn Ting, Devapratim Sarma, Amit Sethi, and Jeffrey Weiss for their help with data collection. Research reported in this publication was supported by the National Institute Of Neurological Disorders And Stroke of the National Institutes of Health under Award Numbers UH3NS107714 and U01NS108922. The content is solely the responsibility of the authors and does not necessarily represent the official views of the National Institutes of Health.

